# Signatures of social pain empathy: general and process-specific brain-wide representations of social exclusion and separation

**DOI:** 10.1101/2024.05.19.594630

**Authors:** Xiaodong Zhang, Peng Qing, Qi Liu, Can Liu, Lei Liu, Xianyang Gan, Kun Fu, Chunmei Lan, Xinqi Zhou, Keith M. Kendrick, Benjamin Becker, Weihua Zhao

## Abstract

Empathy can be elicited by physiological pain, as well as in social contexts. Although physiological and different social contexts induce a strong subjective experience of empathy, the general and context-specific neural representations remain elusive. Here, we combine fMRI with multivariate pattern analysis to establish neurofunctional models for pain empathy triggered by social exclusion and separation. Our findings revealed both overlapping and distinct neural representations for social exclusion and separation empathy across cortical and subcortical regions. This study established an evolutionary model that traces the progression from social pain to physiological pain empathy. In conclusion, this study establishes neural decoding models for pain empathy evoked by social exclusion and social separation, revealing their neural foundations and interconnectedness of empathy induced by social and physiological stimuli. These findings deepen our understanding of the neurobiological mechanisms underlying social pain empathy and provide robust neuromarkers to precisely evaluate empathy across physiological and social domains.

## Introduction

Empathy refers to the ability of individuals to perceive and vicariously experience the experiences of others, enabling them to understand and share their emotional responses, thereby playing a pivotal role in facilitating social interactions ^1,2^. Previous studies have examined empathy in the context of physiological pain ^3,4^, focusing on the anterior insula (AI), the anterior cingulate cortex (ACC), anterior midcingulate cortex (aMCC), and amygdala as components of the typical pain empathy network ^3,5–12^. Yet, in daily life, empathy is often elicited by “social” pain rather than physiological pain. Witnessing others’ social suffering, engaging in empathy towards their situations, and contemplating their thoughts involve additional social processes, such as mentalizing and social functions, which activate regions like the posterior superior temporal sulcus (pSTS) ^13^, the medial prefrontal cortex (mPFC), the precuneus, and the temporoparietal junction (TPJ) ^14–18^. While these regions may serve specific functions in complex social situations ^19^, their precise engagement in the context of social pain empathy remains unclear ^20,21^. Additionally, pain empathy serves as a fundamental form of empathy, whereas social pain empathy is more complex and can manifest in diverse contexts ^22^. The question arises whether social pain empathy is similar across different social contexts, or if it has evolved from basic pain empathy experiences.

Social exclusion and separation induce intense negative affect during social interactions. From an evolutionary perspective, these experiences represent a profound loss of protective social bonds or connections ^23^. Specifically, social exclusion refers to the subjective experience of being rejected during social interactions, such as being ignored or excluded by peers, friends, and others ^24,25^. In contrast, social separation encompasses the state of being physically or emotionally distant ^26^, occurring in scenarios such as the loss of a loved one, romantic breakups, or family separations ^27–29^. Both conditions can elicit a high level of aversive emotional experience, which in turn can induce empathy in individuals observing such situations. While exclusion and separation are conceptually distinct, current research has yet to provide a comprehensive understanding of the neural representations that underlie social pain empathy towards individuals experiencing either of these conditions. The pain empathy triggered by observing noxious stimulation that induces physiological pain can benefit an individual via its evolutionary learning-protective functions ^30^. Conversely, the pain empathy elicited by negative social situations or experiences associated with social pain, such as exclusion or separation, may promote more prosocial helping behaviors ^31–33^. Physiological pain empathy may have evolved as a fundamental form of pain empathy, which was later adapted to more complex social scenarios. Previous studies have identified both overlapping and distinct neural mechanisms involved in physiological and social pain from sensory-discriminate and affective-motivational dimensions ^34–36^. Exploring the neural representations of pain empathy triggered by the same affective dimension but varying social contexts, such as social exclusion and separation, is crucial for advancing our understanding of human complex social interactive behaviors and our empathy towards others emotional states (i.e. distress).

To develop comprehensive and process-specific neurofunctional models that can describe cognitive and affective experiences with large effect sizes ^37^ and at a finer spatial scale ^38^, machine learning-based multivariate pattern classification approaches have been established during recent years and have been successfully applied to pain, pain empathy, and general and specific affective domains ^39–48^. To establish neural representation models of the psychological processes underlying social pain empathy, taking into account contextual and individual variations on the experience level ^49^, we here combined functional magnetic resonance imaging (fMRI) with machine learning-based multivariate pattern analysis (MVPA) ^39,41–43,50^.

To address these issues, in the current study, we deliberately selected natural stimuli (i.e. video clips), as a diversified ecological paradigm, closely resembling real-life social interactions ^51,52^. This was chosen to enhance participants’ engagement and more effectively trigger social pain empathy. Consequently, all stimulus materials in this study were presented as video clips, allowing participants to observe others experiencing either social exclusion or separation from a third-person perspective. This setup allowed us to investigate the neural presentations associated with the common and distinct experiences of social exclusion and separation in social situations.

We employed MVPA to (1) determine specific neural signatures for pain empathy triggered by naturalistic social exclusion stimuli (pain empathy signature of social exclusion, PSSE) depicting individuals experiencing ostracism or being distanced by their peers, and naturalistic social separation stimuli (pain empathy signature of social separation, PSSS) presenting individuals undergoing separations from their family members, romantic partners, and friends separately using a discovery cohort (n=65 healthy participants) and an independent replication cohort (n=35 healthy participants, details see “Stimuli” of the “Methods”; **Figure 1A**); (2) identify the common (PSSEaS) and distinct (PSSEvS) neural signatures underlying these two types of social pain empathy in the discovery cohort and replication cohort; (3) confirm the generalization of developed social pain empathy decoders to other domains including body limbs (n=238) and painful facial expressions (n=238) induced by empathic pain ^46^, as well as the episodic memory retrieval of painful stimuli (n=77, **Figure 1C**); and (4) compare the performance between the established decoders for social pain empathy and other different decoders developed by social pain empathy ^46^(induced by observing painful facial expressions), physical pain empathy ^46^(induced by observing noxious stimulation of others’ body limbs), the combination between social pain empathy and physical pain empathy ^46^, social pain (induced by experienced social rejection^43^), physical pain (induced by heat pain ^43,53^, **Figure 1C**). Together, this systematic examination allows us to establish comprehensive neural models for social pain empathy across varying social contexts and to determine the degree of neurofunctional overlap with the experience of physiological pain, highlighting the translational potential for precisely evaluating empathy across physiological and social domains.

**Fig. 1.**
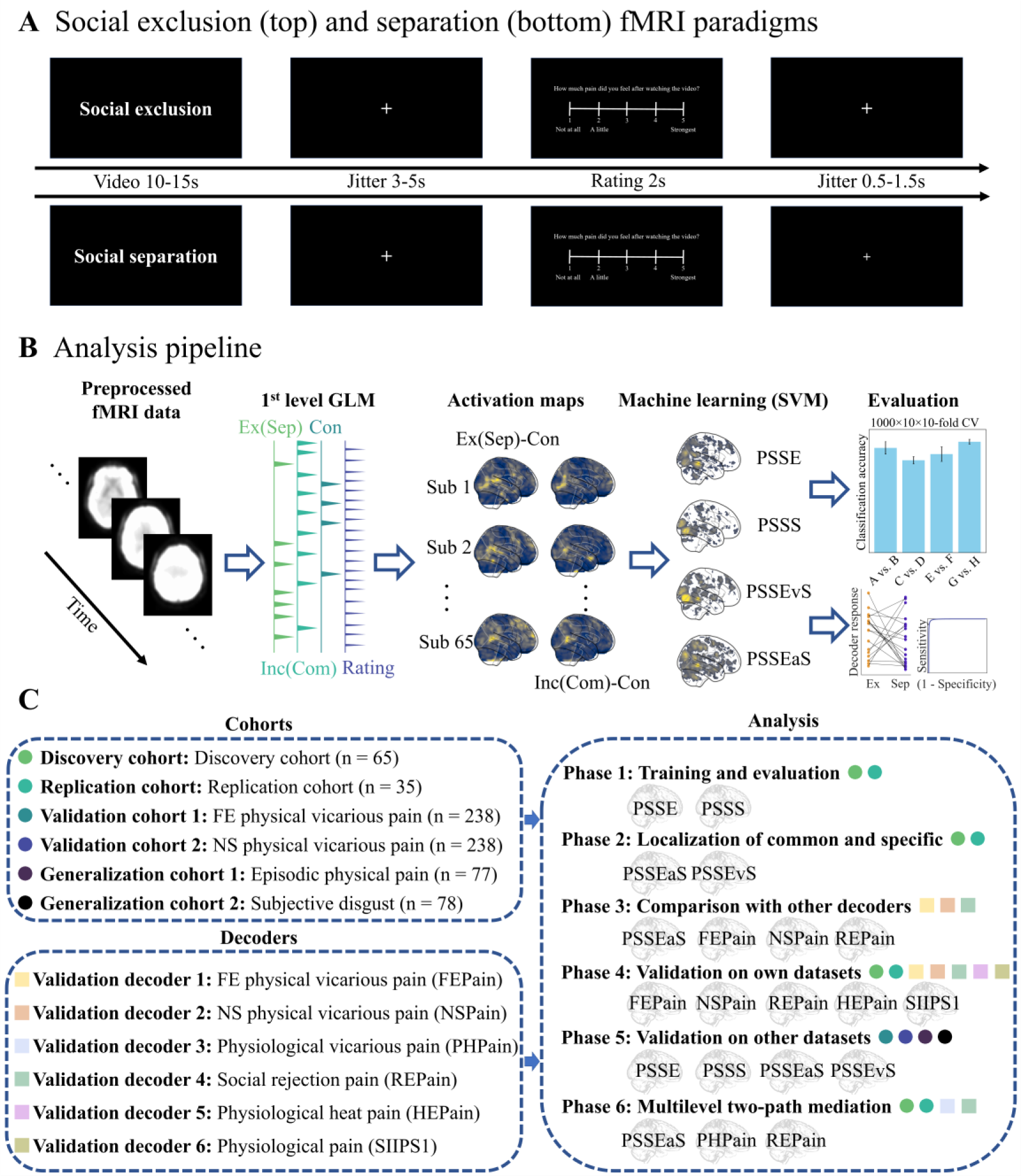
Experimental paradigm, model construction, and analysis workflow. **(A)** Social pain empathy paradigm. The paradigm with a combination with functional magnetic resonance imaging (fMRI) is designed to induce pain empathy through the presentation of social exclusion and social separation video clips. During the exclusion (or separation) session, participants are presented with a total of 60 stimuli, randomly distributed across three fMRI runs. **(B)** Model training and feature selection. Brain activation maps (beta maps) derived from the first-level general linear model (GLM) analysis are extracted as features. Support Vector Machines (SVM) were employed to train whole-brain pain empathy decoders, including PSSE, PSSS, PSSEvS, and PSSEaS, specifically on the discovery cohort (n=65). **(C)** Overview of included datasets, primary analyses, and whole-brain multivariate models. Both discovery and replication cohorts followed the experimental paradigms in Fig. 1A. Validation cohort 1 employs facial pain stimuli to evoke empathic responses (n=238), Validation cohort 2 comprises bodily pain stimuli to evoke empathic responses (n=238), while generalization cohort 1 incorporates a different set of bodily pain image stimuli to recall the content from blurred to clear, to assess the memory performance of empathetic response (n=77), and the generalization cohort 2, consisting of visually evoked subjective disgust (n=78). Primary analyses involve: (1) developing and evaluating PSSE and PSSS models using SVM on the discovery cohort, followed by assessing their performance on both discovery and replication cohorts; (2) exploring shared and distinct voxel-level representations for pain empathy induced by social exclusion and separation stimuli; (3) a comprehensive exploration of decoder performance, comparing it with previous studies on FEPain, NSPain, PHPain, REPain, HEPain and SIIPS1. For a detailed description, please refer to the “Methods” section. Ex, Exclusion; Inc, Inclusion; Sep, Separation; Com, Company. PSSE, Pain empathy signature of social exclusion; PSSS, Pain empathy signature of social separation; PSSEaS, Pain empathy signature of social exclusion and separation; PSSEvS, Pain empathy signature of social exclusion vs. separation; FEPain, Facial Expressions Pain Experience decoder; NSPain, Noxious Stimulation Pain Experience decoder; PHPain, Physical Pain Empathy Signature; REPain, the Social Rejection Experience decoder; HEPain, the Physiological Heat Pain decoder; SIIPS1, the Stimulus Intensity Independent pain signature-1.

## Results

### Behavioral results

A total of 60 video stimuli (22 for exclusion, 22 for inclusion and 16 for control), ranging from 10 to 15 s each, were included for the social exclusion pain empathy task. Inside the scanner, participants were instructed to observe the social experiences of the characters in the videos from a third-person perspective. Following each video, they were prompted to rate their level of pain empathy for the stimulus on a 5-point Likert scale, ranging from ‘not at all’ to ‘strongest’. The design for the social separation pain empathy task mirrored the social exclusion task (for details, see “Methods”). Out of the scanner, they were required to complete arousal and intensity ratings for all stimuli using 5-point Likert scale (1-very low; 5-very high). To compare whether the exclusion and separation stimuli varied by the pain ratings, we conducted paired t tests. Both social exclusion (Ex, M = 2.89, SD = 0.75) and separation (Sep, M = 3.48, SD = 0.73) videos elicited significant negative empathic responses based on self-reported pain ratings, compared to their respective positive (*psFDR* < 0.001) and neutral conditions (*psFDR* < 0.001). However, the pain empathy response induced by the separation was stronger than that induced by the exclusion videos (*t(64)* = 9.47, *pFDR* < 0.001, Cohen’s *d* = 0.80, **Fig. 2A**, more details see Supplementary).

**Fig. 2.**
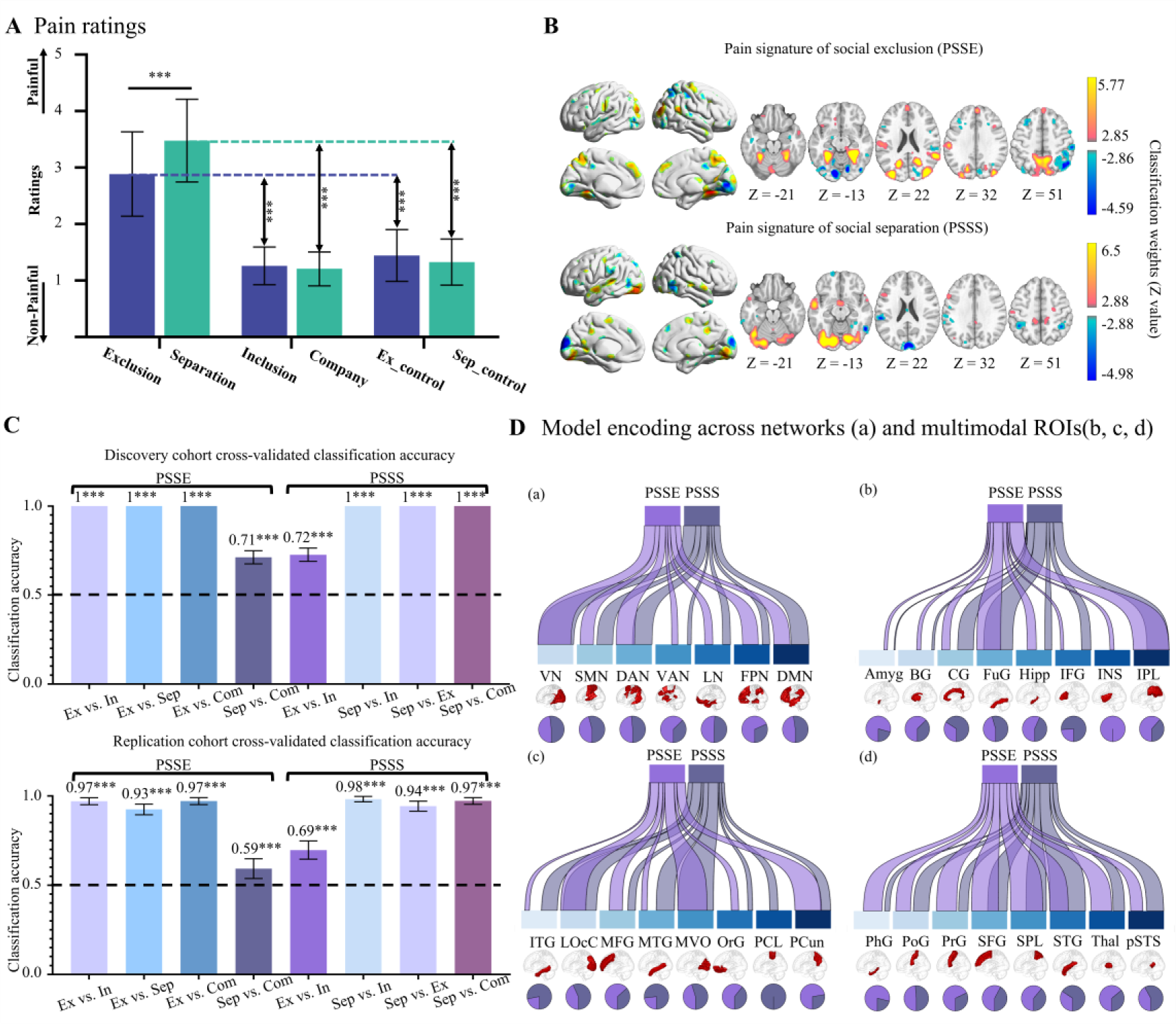
Establishment and performance evaluation of PSSE and PSSS model. **(A)** Behavioral results. Subjective empathy pain ratings for 6 types of stimuli in the discovery cohort (n= 65). Error bars represent standard deviation (SD). **(B)** Core systems for pain empathy to exclusion and separation stimuli. Defined by the conjunction of SVM weight maps and structural coefficients maps. Each model is thresholded at a false discovery rate (FDR) of q<0.05. **(C)** Model performance evaluation. Two-choice classification accuracy evaluated using 1000 iterations of 10×10-fold cross-validation on discovery (n=65) and replication (n=35) cohorts. Chance level is indicated by the dashed line, and error bars represent the standard deviation of 1000 results. **(D)** PSSE and PSSS streamgraphs. Representing the spatial similarity computed as cosine similarity of the two core systems across 7 networks and 24 brain regions. Ribbons are normalized to the maximum cosine similarity across all networks and ROIs. Pie charts display the relative contribution of each model to each network or ROI region (i.e., the percentage of voxels with the highest cosine similarity for each predictive map). ***P<0.001, all p values are FDR-corrected. Ex, Exclusion; Inc, Inclusion; Sep, Separation; Com, Company. Ex_control, the control condition for exclusion session; Sep_control, the control condition for separation session; VN, Visual Network; SMN, Somatomotor Network; DAN, Dorsal Attention Network; VAN, Ventral Attention Network; LN, Limbic Network; FPN, Frontoparietal Network; DMN, Default Network; Amyg, Amygdala; BG, Basal Ganglia; CG, Cingulate Gyrus; FuG, Fusiform Gyrus; Hipp, Hippocampus; IFG, Inferior Frontal Gyrus; INS, Insula; IPL, Inferior Parietal Lobule; ITG, Inferior Temporal Gyrus; LOcC, Lateral Occipital Cortex; MFG, Middle Frontal Gyrus; MTG, Middle Temporal Gyrus; MVO, MedioVentral Occipital Cortex; OrG, Orbital Gyrus; PCL, Paracentral Lobule; PCun, Precuneus; PhG, Parahippocampal Gyrus; PoG, Postcentral Gyrus; PrG, Precentral Gyrus; SFG, Superior Frontal Gyrus; SPL, Superior Parietal Lobule; STG, Superior Temporal Gyrus; Thal, Thalamus; pSTS, posterior Superior Temporal Sulcus.

### Identifying stimulus-type-specific brain models

Exclusively within the gray matter mask ^54^, a multivariate machine learning pattern analysis was conducted throughout the entire brain ^39^. The pain empathy signature of social exclusion (PSSE) or social separation (PSSS) is presented with voxel classification weights across whole-brain, separately. These weights were determined with a threshold of P < 0.05 (FDR corrected), based on bootstrap tests with 10,000 iterations. The reliable weights of significant regions were effectively used to distinguish exclusion from other conditions. The retained threshold is strictly for visualization purposes, and all voxel weights were employed in the classification. The results indicate that empathic processes are characterized by significant classification weights across a distributed network of subcortical and cortical systems (**Fig. 2B**). Specifically, pain empathy induced by the naturalistic exclusion involved core empathy-related networks including the left anterior insula, dorsal anterior cingulate cortex (dACC), and social processing network, such as left posterior cingulate gyrus, thalamus, posterior superior temporal sulcus (pSTS), medial prefrontal cortex (mPFC), precuneus (PCun) and TPJ. On the other hand, pain empathy triggered by separation activated common (i.e. dACC, pSTS, mPFC, PCun, TPJ) and specific brain regions, such as ventrolateral prefrontal cortex (vlPFC), supplementary motor area (SMA), supra marginal gyrus, left precentral gyrus and left hippocampus (**Fig. 2B**).

### Benchmarking and generalization of PSSE and PSSS performance

A robust evaluation of the performance of the PSSE and PSSS models in the discovery cohort was conducted via 1000 iterations of a 10×10-fold cross-validation procedure (details **see Methods**). The PSSE model demonstrated a remarkable capacity to discriminate between various types of stimuli (Ex vs. other three conditions, accuracy 100%; Sep vs. Com, 71.2 ± 3.7%; **Fig. 2C**). Additionally, the PSSS model exhibited significant classification performance between separation and others: Sep vs. other three conditions, 100%, as well as Ex vs. Inc, 72.7 ±3.7% (**Fig. 2C**).

To validate the generalizability of our models, it is imperative to establish the stability of group-level neural features across different cohorts. The discrimination performance of the PSSE and PSSS models in the replication cohort was as follows: for the PSSE model: Ex vs. Inc, 97.0 ± 2.0%; Ex vs. Sep, 92.5 ± 3.0%; Ex vs. Com, 97.1 ± 2.0%; Sep vs. Com, 59.3 ±5.6%; **Fig. 2C**) and for the PSSS model: Sep vs. Com, 97.2 ±1.9%; Sep vs. Ex, 94.3 ±2.8%; Sep vs. Inc, 98.3 ±1.5%; Ex vs. Inc, 69.0 ±5.1% (see **Fig. 2C**). These findings demonstrate that the developed decoders showed consistent and reliable performance across two cohorts.

### Individual model contributions to social pain empathy

Based on the computed classification accuracies and overlapping brain regions observed in these two models, we hypothesized that they might be intrinsically connected. To ascertain the specific and shared neural representations, we employed the Human Brainnetome atlas to divide the brain into seven networks and 24 regions of interest (ROIs). By calculating the spatial similarity (cosine similarity) of the two models in each region, we estimated the relative contributions of each network or ROI to each model. Both models exhibited stable predictive voxels across all seven networks, yet the relative contributions of individual networks (excluding the visual network, as both empathic responses were elicited using video stimuli) differed between the models. Specifically, the ventral attention network (VAN) and frontoparietal network (FPN) exhibited stronger contributions to PSSE than to PSSS, whereas the somatomotor network (SMN), dorsal attention network (DAN), visual network (VN), and default network (DMN) displayed stronger contributions to PSSS (**Fig. 2D**). Similarly, an analysis of the 24 ROIs revealed that while representations of PSSE and PSSS were relatively similar in corresponding areas of the visual network (i.e. LOcC and MVOoC), notable differences in contributions were observed in other brain regions between the two models (**Fig. 2D**). These findings provide insights into the neural representations underlying the distinct empathic responses elicited by social exclusion and separation contexts (details see **Supplementary Table. 1**).

### Commonalities and distinctions between social exclusion and social separation

To further investigate the neural mechanisms underlying pain empathy for social exclusion and separation, we developed specific neural decoders, analogous to PSSE and PSSS, based on the discovery cohort. Specifically, we developed PSSEaS (Pain Signature for Social Exclusion and Separation), aimed at discriminating between exclusion and separation vs. inclusion and company (with 1 indicating exclusion and separation, and -1 indicating inclusion and company), and PSSEvS (Pain Signature for Social Exclusion vs. Separation), designed to distinguish between exclusion and separation (with labels 1 representing exclusion and -1 representing separation). **Figure 3A** depicts the threshold maps of PSSEaS and PSSEvS (q<0.05, FDR correction, 10000 iterations of bootstrap). PSSEaS offers a more distributed and extensive network of brain regions that collectively contribute to pain empathy in both exclusion and separation context. These regions, such as ACC, insula, TPJ, and pSTS, have been associated with pain empathy and are activated during both social exclusion and separation stimuli. On the other hand, PSSEvS reveals the core regions involved in distinguishing exclusion from separation. For instance, regions such as IPL, thalamus, ITG, and PCun exhibited positive correlations with the exclusion condition, while regions like dACC, MFG, INS, and portions of the occipital lobe showed positive correlations with the separation condition.

**Fig. 3.**
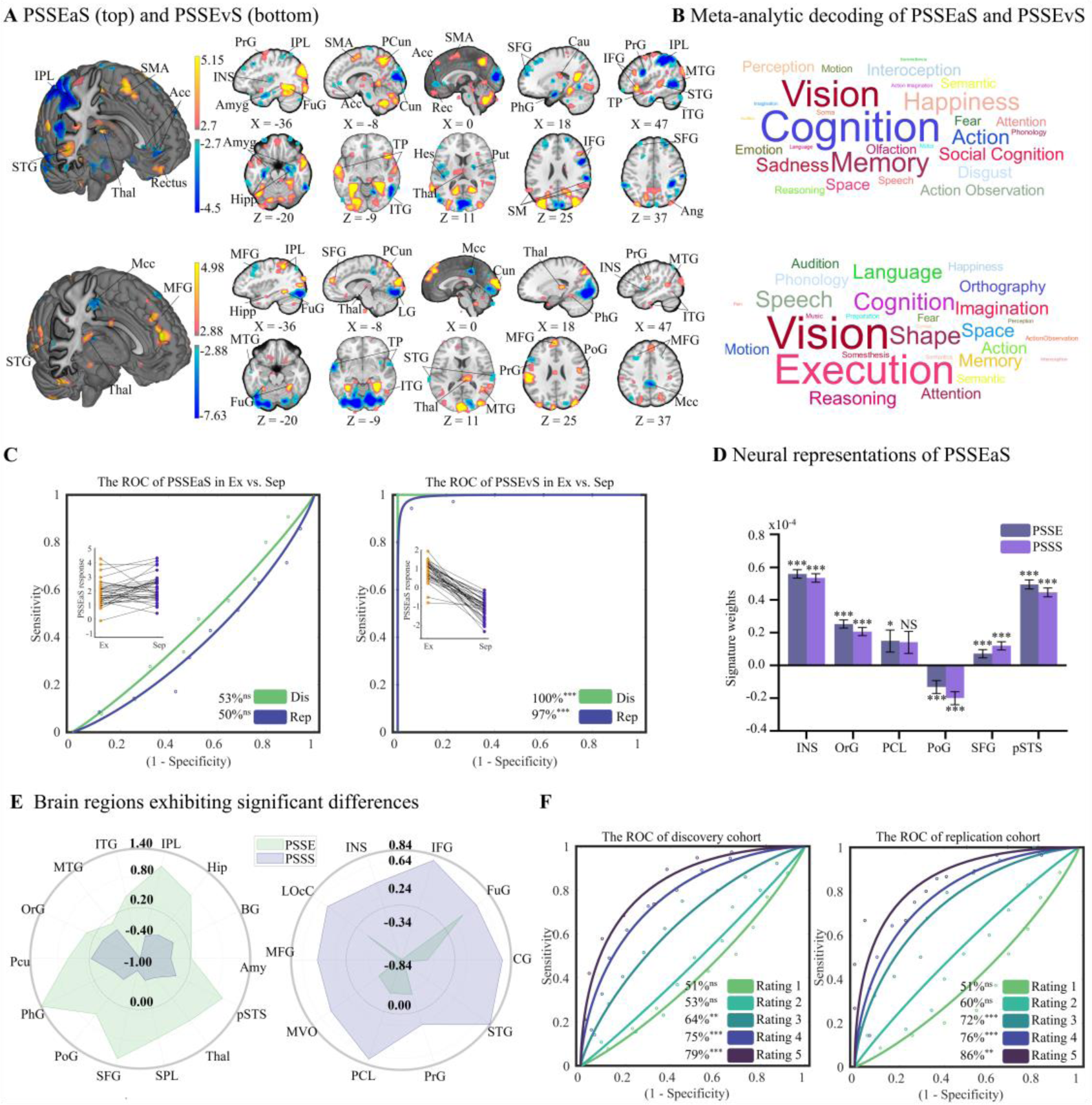
Neural level disparities and commonalities of pain empathy to social exclusion and separation stimuli. **(A)** Stimuli-specific and common neural core systems underlying pain empathy to social exclusion and separation stimuli. **(B)** Word cloud of PSSEaS and PSSEvS. Visualizing the functional behavioural terms of the top 20 atlas regions from meta-analytic decoding (provided by the Brainnetome website), highlighting the distinctiveness of social exclusion and separation pain empathy patterns. Font size reflects the strength of relevance. **(C)** Performance evaluation of PSSEvS and PSSEaS. Assessing the discriminatory capacity of the models in distinguishing exclusion and separation stimuli across discovery and replication cohorts. **(D)** Similar neural representations of pain empathy of exclusion and separation. Average SVM weights within contiguous regions, based on the core regions of PSSEaS. **(E)** Differential neural representations of pain empathy of exclusion and separation. For display purposes, weight values have been multiplied by 10,000. Average SVM weights computed within contiguous regions, based on the core regions of PSSEvS. **(F)** Classification accuracy of different empathy pain scores for exclusion and separation. * P<0.05, ** P<0.01, *** P<0.001, NS, not significant, all p values are FDR-corrected. Ex, Exclusion; Inc, Inclusion; Sep, Separation; Com, Company. Dis, Discovery; Rep, Replication. Amyg, Amygdala; BG, Basal Ganglia; CG, Cingulate Gyrus; FuG, Fusiform Gyrus; Hipp, Hippocampus; IFG, Inferior Frontal Gyrus; INS, Insula; IPL, Inferior Parietal Lobule; ITG, Inferior Temporal Gyrus; LOcC, Lateral Occipital Cortex; MFG, Middle Frontal Gyrus; MTG, Middle Temporal Gyrus; MVO, MedioVentral Occipital Cortex; OrG, Orbital Gyrus; PCL, Paracentral Lobule; PCun, Precuneus; PhG, Parahippocampal Gyrus; PoG, Postcentral Gyrus; PrG, Precentral Gyrus; SFG, Superior Frontal Gyrus; SPL, Superior Parietal Lobule; STG, Superior Temporal Gyrus; Thal, Thalamus; pSTS, posterior Superior Temporal Sulcus; Mcc, Middle Cingulate Cortex; LG, Lingual; Cun, Cuneus; Acc, Anterior Cingulate Cortex; TP, Temporal Pole; SMA, Supplementary Motor Area; Rec, Rectus; SM, Supramarginal Gyrus; Ang, Angular Gyrus; Put, Putamen; Cau, Caudata; Hes, Heschl.

The performance of both PSSEaS and PSSEvS was thoroughly evaluated in both the discovery and replication cohorts. PSSEaS failed to differentiate between social exclusion and separation in either cohort (discovery: 53% ± 4.4%, p = 0.54, AUC = 0.43; replication: 50% ±6.0%, p = 1, AUC = 0.38), indicating a lack of discrimination between two types of pain empathy. By contrast, PSSEvS accurately predicted pain empathy evoked by exclusion or separation (discovery: 100%, p < 0.001, AUC = 1.00, and Cohen’s d = 7.99; replication: 97% ± 2.0%, p < 0.001, AUC = 0.99, and Cohen’s d = 3.65, **Figure 3C**).

We decoded the top 20 atlas regions that overlapped with the thresholded PSSEaS and PSSEvS maps. Utilizing the functional behavioral decoding provided by the Brainnetome website, we mapped these regions to word clouds. The font size within the word cloud is directly proportional to the activation likelihood ratio assigned to each term by BrainMap, with larger ratios corresponding to larger font sizes. For PSSEaS, the meta-analysis revealed involvement in cognition, vision, memory, social cognition, and a range of emotions. In contrast, PSSEvS decoded terms related more closely to cognitive and perception functions, such as execution and vision etc. (**Figure 3B**).

### Neural representations between exclusion and separation

To further validate the core contributing regions of PSSEaS, we categorized these regions into 24 distinct yet standardized areas based on the Human Brainnetome atlas in order to explore the common neural representations of exclusion and separation pain empathy. As depicted in **Figure 3D**, similar patterns of classification weights between PSSE and PSSS were observed in the anterior insula, orbital gyrus (OrG), paracentral lobule (PCL), postcentral gyrus (PoG), superior frontal gyrus (SFG), and pSTS. Particularly, the classification weights in the PCL region were not significant in PSSS (i.e., PCL did not contribute significantly to separation empathy using one-sample t-test).

Similarly, to further explore the neural patterns specific to exclusion and separation pain empathy, we categorized the core system of PSSEvS into 24 subregions and computed the classification weights of these regions in both PSSE and PSSS. We found that activation in 14 regions (Amygdala-Amyg, Basal Ganglia-BG, Hippocampus-Hipp, Inferior Parietal Lobule-IPL, Inferior Temporal Gyrus-ITG, Middle Temporal Gyrus-MTG, Orbital Gyrus-OrG, Precuneus-PCun, Parahippocampal Gyrus-PhG, Postcentral Gyrus-PoG, Superior Frontal Gyrus-SFG, Superior Parietal Lobule-SPL, Thalamus-Tha, Posterior Superior Temporal Sulcus-pSTS, **Figure 3E**) was more strongly predictive of pain empathy evoked by exclusion. Additionally, activation in 10 brain regions (Cingulate Gyrus-CG, Fusiform Gyrus-FuG, Inferior Frontal Gyrus-IFG, Insula-INS, Lateral Occipital Cortex-LOcC, Middle Frontal Gyrus-MFG, MedioVentral Occipital Cortex-MVOcC, Paracentral Lobule-PCL, Precentral Gyrus-PrG, Superior Temporal Gyrus-STG; **Figure 3E**) was more strongly predictive of pain empathy evoked by separation. The significant weight differences in these regions were primarily driven by significant negative weights for pain empathy induced by both exclusion and separation.

### Exploring the discriminative ability of empathy strength across multiple levels

Returning to an intriguing topic, our initial stimulus screening behavioral experiment and subsequent fMRI experiment consistently revealed higher self-reported pain intensity under social separation than social exclusion stimuli. This prompts a question in the present study whether the observed differences between social exclusion and separation stem from the induction conditions themselves or from the varying intensity of empathy experienced under these conditions, ultimately leading to distinct neural representations.

To address this question, we constructed a new GLM model, incorporating participants’ subjective ratings (**see Methods**). Using the PSSEvS classifier, specially tailored for distinguishing between exclusion and separation, we compared the two conditions at equivalent rating levels (e.g., exclusion-rating 1 vs. separation-rating 1, exclusion-rating 5 vs. separation-rating 5). Initial findings from the discovery cohort suggested that the classifier was unable to differentiate between exclusion and separation at rating 1 (51 ± 4.4 % accuracy, *pFDR* = 0.93) and rating 2 (53 ± 4.4% accuracy, *pFDR* = 0.54). However, as the subjective ratings of the stimulus materials increased among participants, we observed a gradual improvement in classification accuracy. This trend was evidenced by rating 3 (64 ±4.2 % accuracy, *pFDR* = 0.002, Cohen’s *d* = 0.55), rating 4 (75 ± 4.0 % accuracy, *pFDR* < 0.001, Cohen’s *d* = 1.06), and rating 5 (79 ± 4.3% accuracy, *pFDR* < 0.001, Cohen’s *d* = 1.39; **Figure 3F**).

Consistent with the findings from the discovery cohort, the replication cohort also exhibited similar results: rating 1 (51 ± 6.0 % accuracy, *pFDR* = 0.91), rating 2 (60 ±5.9 % accuracy, *pFDR* = 0.12), rating 3 (72 ±5.4 % accuracy, *pFDR* < 0.001, Cohen’s *d* = 0.91), rating 4 (76 ± 5.6 % accuracy, *pFDR* < 0.001, Cohen’s *d* = 1.15), and rating 5 (86 ± 5.6 % accuracy, *pFDR* < 0.001, Cohen’s *d* = 1.56; **Figure 3F**). These consistent patterns across cohorts suggest that the neural representations associated with the two conditions (exclusion vs. separation) are primarily attributed to the specific types of pain empathy stimuli. Furthermore, it appears that the intensity of pain empathy elicited by the stimulus does not play a significant role in differentiating the neural representations of social exclusion and separation.

### Relationship between social pain empathy and physiological pain empathy

Previous studies have shown shared neural patterns between social pain empathy (i.e. social exclusion) and physiological pain empathy ^43^. However, this study aims to expand our understanding of the interrelationship between different stimuli-specific types of social pain empathy (exclusion and separation), and their connections with pain empathy for physiological pain, separately. Here, we undertook a comparative analysis of the models generated in this study with those derived from visual-induced physiological pain empathy datasets (validation cohort 1 and validation cohort 2) ^43^, physiological pain empathy induced by episodic memory (generalization cohort 1), visual-induced subjective disgust dataset (generalization cohort 2)^40^ and previously established models including social pain empathy ^46^(induced by painful facial expressions, FEPain), physiological pain empathy ^46^(induced by observing noxious stimulation of others’ body limbs, NSPain), the combination between social pain empathy and physiological pain empathy (PHPain), social pain ^43^(induced by experienced social rejection, REPain), physical pain (induced by heat pain, HEPain ^43^ and SIIPS1 ^53^, **Figure 1C**, more details see “Extended exploration of decoder performance” section).

First, we used term-based meta-analyses in Neurosynth to generate a whole-brain region of interest on “Empathy” (**Supplementary Figure. 4**) and calculated the spatial similarity (cosine similarity) of the three decoders related to empathy: PSSEaS, FEPain and NSPain. Overall, PSSEaS is more widely distributed than FEPain and NSPain (**Figure 4A, Supplementary Table. 2**). In particular, in INS, physiological pain empathy (FEPain, NSPain) was stronger than social pain empathy (PSSEaS). In addition, we plotted decoder weight patterns for the models (including REPain) on INS, cerebellum, Periaqueductal Gray (PAG), PCun, dorsomedial prefrontal cortex (dmPFC), and Rectus. The results showed that social pain empathy and physiological pain were similar but different in empathy processing areas, such as INS, Cereb and PCun. The neural representations of social pain empathy and social pain also show similarities, with significant correlations in PCun, PAG, dmPFC and Rectus (**Figure 4B, Supplementary Table. 3**).

**Fig. 4.**
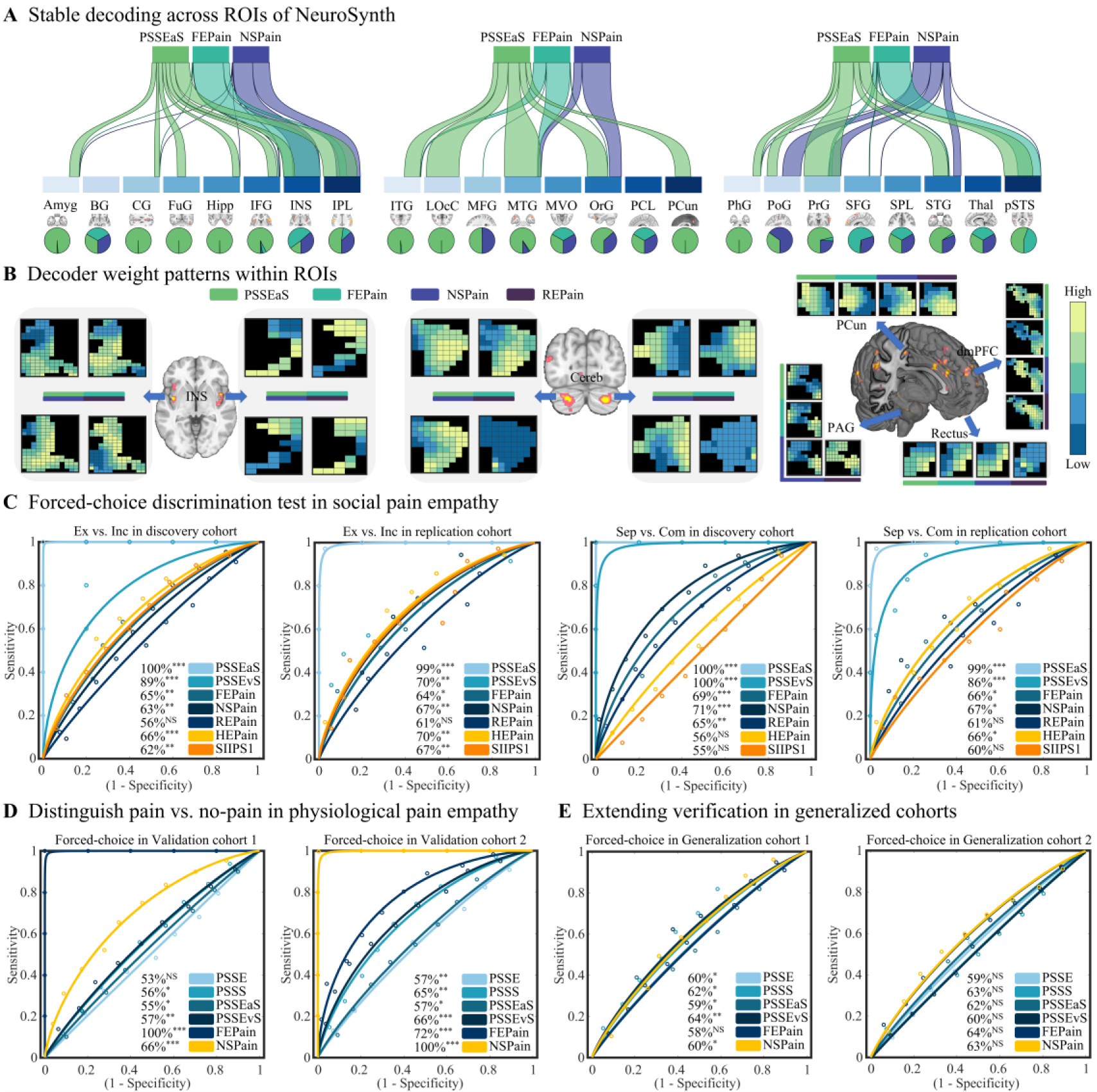
Extended exploration of decoder performance. **(A)** The spatial similarity (cosine similarity) between stable decoding maps and selected ROIs (from NeuroSynth search “Empathy”). **(B)** Supplementary to **(A)**, the weights of PSSEaS, FEPain, NSPain, and REPain in multiple empathic brain regions are shown. **(C)** Distinguishing exclusion vs. inclusion and separation vs. company using PSSEaS, PSSEvS, FEPain, NSPain, REPain, HEPain and SIIPS1. **(D)** Distinguishing pain vs. no-pain in physiological pain empathy using PSSE, PSSS, PSSEaS, PSSEvS, FEPain, and NSPain. **(E)** Extending verification in generalized cohorts using PSSE, PSSS, PSSEaS, PSSEvS, FEPain, and NSPain. * P<0.05, ** P<0.01, *** P<0.001, NS, not significant, all p values are FDR-corrected. Amyg, Amygdala; BG, Basal Ganglia; CG, Cingulate Gyrus; FuG, Fusiform Gyrus; Hipp, Hippocampus; IFG, Inferior Frontal Gyrus; INS, Insula; IPL, Inferior Parietal Lobule; ITG, Inferior Temporal Gyrus; LOcC, Lateral Occipital Cortex; MFG, Middle Frontal Gyrus; MTG, Middle Temporal Gyrus; MVO, MedioVentral Occipital Cortex; OrG, Orbital Gyrus; PCL, Paracentral Lobule; PCun, Precuneus; PhG, Parahippocampal Gyrus; PoG, Postcentral Gyrus; PrG, Precentral Gyrus; SFG, Superior Frontal Gyrus; SPL, Superior Parietal Lobule; STG, Superior Temporal Gyrus; Thal, Thalamus; pSTS, posterior Superior Temporal Sulcus; Cereb, Cerebellum; PAG, Periaqueductal Gray; Rec, Rectus; dmPFC, Dorsomedial Prefrontal Cortex.

In the discovery cohort, the REPain model successfully differentiated between separation and company stimuli (65 ±4.2 % accuracy, *pFDR* < 0.01, Cohen’s *d* = 0.57) but failed between exclusion and inclusion conditions (56 ±4.4% accuracy, *pFDR* = 0.19; **Figure 4C**). In the replication cohort, REPain failed to differentiate any of the conditions. Given that the REPain model was trained using the data from participants observing their ex-partners, these results further emphasized the specificity of our exclusion and separation.

When NSPain, and FEPain were utilized in both discovery and replication cohorts to distinguish between negative and positive conditions, both classifiers, which are responsible for physiological pain empathy, demonstrated a significant capacity to distinguish social pain empathy. Forced-choice discrimination test on Exclusion and Inclusion, HEPain and SIIPS1 showed significant ability, but could not significantly distinguish between Separation and Company (**Figure 4C**; Statistical information is provided in **Supplementary Table. 4**). Additionally, when PSSE, PSSS, PSSEaS, PSSEvS, FEPain, and NSPain were applied to validation cohort 1, PSSS, PSSEaS, and PSSEvS were able to significantly differentiate physiological pain empathy. For the validation cohort 2, all decoders significantly recognized physiological pain empathy (**Figure 4D; Supplementary Table. 5**).

In additional analysis, we assessed the classification performance of pain empathy models for episodic memory-induced physiologic pain empathy and visually induced negative emotion (i.e. disgust). In the generalization cohort 1, we found that four models were able to significantly identify pain and non-pain conditions, while FEPain did not, likely due to the specific focus on limb pain empathy in the cohort. In the generalization cohort 2, all pain empathy decoders failed to significantly classify subjective disgust emotions (**Figure 4D; Supplementary Table. 5**).

Furthermore, a multi-level mediation model was adopted to explore the neural representations encoding subjective pain empathy experiences in PSSEaS, PHPain, REPain. In both the discovery and replication cohorts, we found that the PHPain (the combination of physical pain empathy induced by observing noxious stimulation of others’ body limbs and social pain empathy induced by observing painful expressions) response partially mediated the effect of PSSEaS response on ratings of social pain empathy (**Figure 5A**). Similarly, the results also showed that PSSEaS response partially mediated the effect of PHPain response on social pain empathy ratings (**Figure 5B**). However, the REPain response failed to mediate the effect of PSSEaS response on social pain empathy ratings (**Figure 5C**), and PSSEaS response completely mediates the association between REPain response and subjective social pain empathy ratings (**Figure 5D**). Thus, it is plausible to hypothesize an inherent neural connection between social pain empathy and physiological pain empathy ^34,35^.

**Fig. 5.**
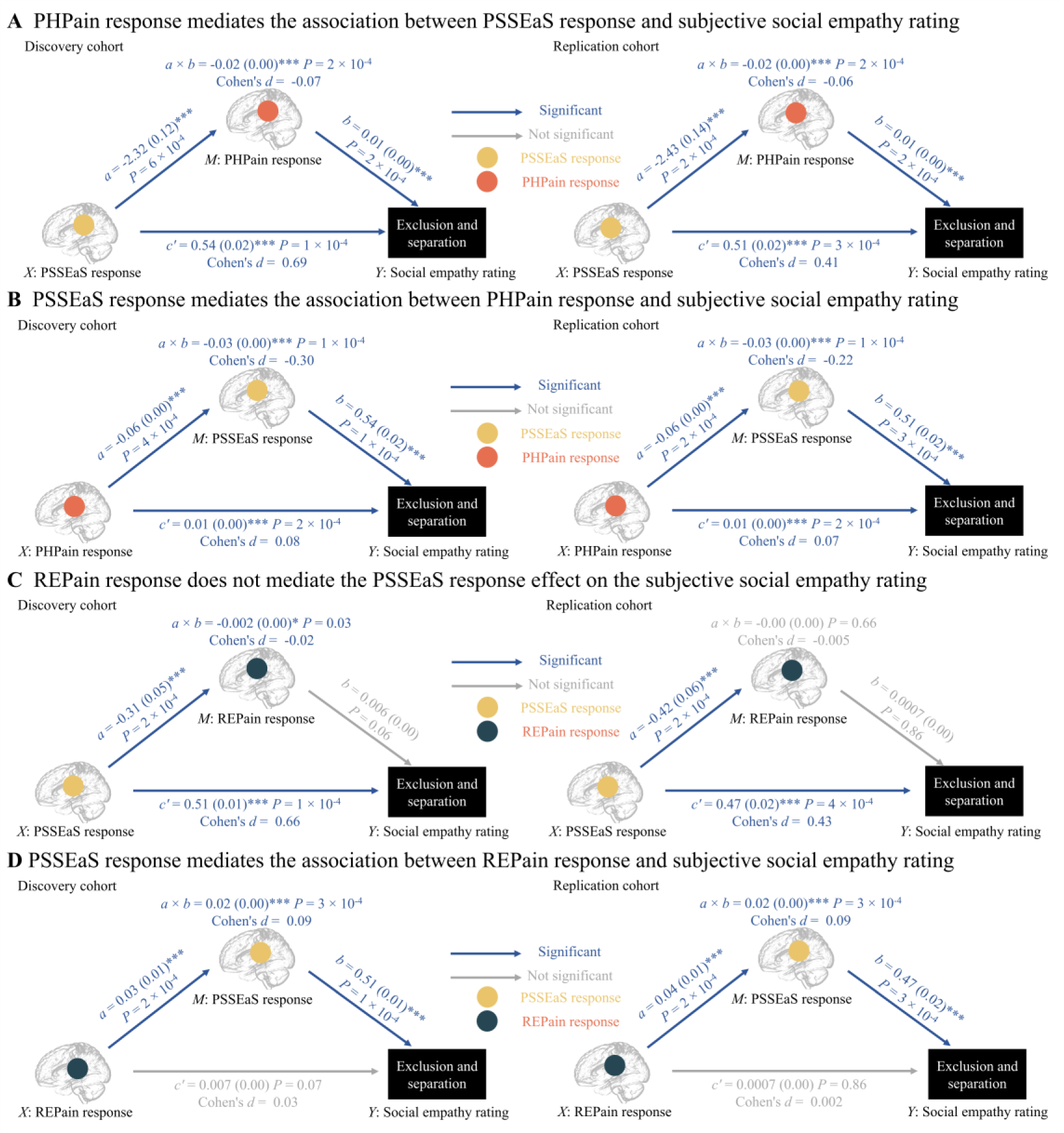
Separable characteristics of social pain empathy, physical pain empathy and social pain. **(A)** The physiological pain empathy model (PHPain response, a combined model of FEPain and NSPain) is examined for its mediating role in the relationship between PSSEaS response and subjective social empathy ratings in both discovery and replication cohorts. **(B)** The mediating role of the social pain empathy model (PSSEaS response) in the connection between PHPain response and subjective social empathy ratings. **(C)** Non-mediation of subjective social empathy ratings by REPain response in the effect of PSSEaS response in both the discovery and replication cohorts, regardless of the social condition of exclusion or separation. **(D)** The PSSEaS response serves as a mediator in linking the REPain response to subjective social empathy ratings. * P<0.05, ** P<0.01, *** P<0.001, NS, not significant, all p values are FDR-corrected.

## Discussion

Previous animal models suggested that shared emotion (i.e. emotional contagion involved in pain empathy) promotes prosocial behavior^55^ and understanding how empathy for social pain is encoded in the brain remains a central question in cognitive and interpersonal neuroscience. Social pain induced by social exclusion shares a common neural basis with physiological pain, yet stimulus-specific variations in empathy towards social pain are observed across distributed brain regions. This study provided evidence for both common and stimulus-specific neural representations of empathy for social pain triggered by social exclusion and separation, leveraging machine learning-based neural decoding. Subjective empathy for social pain evoked by both social exclusion (PSSE) and social separation (PSSS) stimuli was encoded in distributed subcortical and cortical brain systems including general empathy (i.e. dACC) and social processing networks (i.e. pSTS, mPFC, PCun, TPJ). Furthermore, the model for empathy towards social pain (PSSEaS) generalized to the processing of physiological pain empathy as well as the retrieval of memories related to physiological pain. Notably, other developed physiological pain decoders showed limited sensitivity to social pain empathy. Furthermore, the neural signature of vicarious physiological pain empathy induced by observation of noxious stimulation of others’ body limbs and painful facial expressions (PHPain) mediated the response of the model we developed (PSSEaS) on subjective pain ratings to social exclusion and separation but not the rejection experienced decoder (REPain), suggesting an evolutionary perspective from basic pain empathy to social pain empathy involved in more complex social experiences. However, the neural representations of stimulus-specific social pain empathy appeared to be distinct and separated, with the model engaging regions associated with mentalizing and social networks for pain empathy triggered by social exclusion, whereas it involved regions related to pain perception and face processing for the pain empathy induced by social separation. Collectively, the current study offers a comprehensive neural signature of stimulus-specific social pain empathy and enabling more robust and generalizable predictions of subjective social pain.

Previous studies emphasized the significance of core regions, such as the bilateral anterior insula, anterior cingulate and midcingulate cortex, in pain empathy, which are often also observed during the experience of nociceptive stimulation. However, we combined a multimodal experimental design and multivariate pattern decoders ^45,54^ for social pain empathy to uncover whether neural representations of social pain empathy vary across stimulus types or a combination of both. This approach allowed us to integrate neural responses from distributed brain regions and numerous voxels, overcoming the limitations of traditional univariate analysis methods. Notably, the induction of social pain empathy, such as watching others being socially excluded or experiencing a separation, is a remarkably intricate and complex process that elicits a broad range of emotional and cognitive responses ^23,56^. Our findings indicated that pain empathy evoked by social exclusion and separation engaged a diverse and widely distributed set of neural representations. Notably, subcortical regions contributed significantly to mentalizing processes, such as the thalamus, putamen, hippocampus, while cortical regions involved in social interaction and somatosensory processing (i.e. ACC, IFG, insula, IPL, precuneus, STG). Thus, we were able to move beyond the challenges of inferring the “pain empathy” state solely based on brain activity ^57^, capturing the collective activity of distributed brain regions across the entire brain ^4^, and addressing issues related to insufficient effect size for brain activation regions ^37,58^.

Furthermore, naturalistic exclusion stimuli elicited pain empathy, involving core empathy-related networks including left anterior insula, dACC, as well as social processing networks, such as left posterior cingulate gyrus, thalamus, pSTS, mPFC, precuneus and TPJ. Conversely, separation stimuli triggered pain empathy and activated common (i.e. dACC, pSTS, mPFC, precuneus, TPJ) and specific brain regions, such as vlPFC, SMA, supramarginal gyrus, left precentral gyrus and left hippocampus. The posterior superior temporal sulcus (pSTS) is engaged in representing others’ belief states ^6^, while the mPFC is responsible for attributing and storing enduring mental states^59^. The precuneus contributes to mental imagery, and TPJ is involved in inferring transient mental states ^17^. Collectively, these regions are more associated with higher-order social cognitive abilities ^60^, highlighting their significance in understanding and interpreting social context. Additionally, a contrast analysis of the pain empathy models for exclusion and separation revealed engagement in regions such as MCC, precuneus, insula, STG. These findings suggest that common and distinct neural networks are activated when individuals experience empathy towards exclusion and separation, further underscoring the complexity of social pain empathy processing.

Although no single network was responsible for predicting subjective pain ratings across both social exclusion and separation conditions, some networks demonstrated stronger contributions to pain empathy under specific conditions. Specifically, the somatomotor network, dorsal attention network, visual network, and default network were more involved in pain empathy during social separation. In contrast, the ventral attention and frontoparietal networks exhibited stronger contributions to pain empathy by social exclusion. From the ROI perspective, specific regions such as amygdala, pSTS, precuneus, IPL and hippocampus exhibited greater contributions for pain empathy under social exclusion ^20,61^, while other regions including IFG, INS, MFG were more engaged in pain empathy during social separation. These findings suggested a process-specific and distinct adaption to another’s sensory and affective state, potentially resulting from the different physiological distances created by social exclusion (perceived as more distant) and social separation (perceived as closer).

We next determined if the observed disparities between social exclusion and separation originated solely from the distinct induction conditions or varied pain empathy intensity ratings, ultimately resulting in distinct neural representations. Our behavioral findings indicated that social separation stimuli evoked higher pain ratings compared to exclusion stimuli. This is consistent with the understanding that social separation, akin to the loss of a loved one, is among the most devastating forms of social pain, often triggering intense psychological distress^23^. Notably, these distinct neural representations appear to be primarily process-specific, rather than being solely determined by different levels of pain ratings, further reinforcing our results. The experiences of social pain were characterized by an unpleasant or negative feeling associated with actual or potential harm to one’s sense of social connection or social value, such as social rejection, exclusion or loss ^23^. Specially, social exclusion pertains to the subjective sense of being ignored or rejected during social interactions ^24,25^, whereas social separation encompasses a state of physiological or emotional distance^26^. These two situations of social disconnection elicited different social pain experiences with the former more associated with the devaluation of self but not the latter ^23^, ultimately leading to process- or stimulus-specific neural representations. Additionally, we also conducted a comprehensive comparison between the validated decoders ^43,46^ and our own decoders to evaluate their performance in decoding social pain empathy ranging from social exclusion to social loss, as well as physiological pain empathy. Our decoders consistently demonstrated that social pain empathy can predict physiological pain responses, but not vice versa. Combined with the multilevel two-path mediation analysis, these findings suggest that while regions involved in physiological pain empathy may also be engaged in social pain empathy, the latter involves a more intricate blend of cognitive and affective responses.

There are several limitations in the current study. Firstly, while the social pain empathy decoders have been examined in the social context of social exclusion and separation, we suggest extending the generalization to some other diverse contexts, such as negative evaluation, where individuals face critical judgments or disapproval from others, such as a negative social evaluation context although both negative evaluation and social exclusion are instances of the devaluation of self^62^. Additionally, the current MVPA results reflect neural activity and distributed activation-based pattern rather than functional connectivity features or functional connectivity-based patterns ^63,64^, and thus requires some caution in interpretation.

In conclusion, the present study compared the corresponding whole-brain neural signatures triggered by empathy for social pain experienced through social exclusion and separation. These visually induced patterns of social pain empathy were validated and generalized across participants, paradigms and study contexts. Our findings suggested that social pain empathy was encoded in multiple distributed brain regions or networks rather than isolated brain areas. Specifically, social pain empathy induced by social exclusion and separation exhibited both common and distinct neural patterns. Notably, their patterns were not solely determined by different levels of pain ratings, but rather reflected stimulus-specific processes. Overall, the study contributed to our understanding of the neurobiological mechanism underlying pain empathy. Dysregulations in social pain empathy are implicated in various mental disorders (i.e. social anxiety ^65^), suggesting that process-specific neural signatures could potentially serve as MVPA-based markers for precise diagnosis.

## Materials and Methods

### Participants in the discovery and replication cohort

The discovery cohort consisted of 65 healthy participants (34 females; mean ± SD age = 20.78 ± 1.62 years) who were recruited from the University of Electronic Science and Technology of China (UESTC). All participants underwent screening to ensure that all were right-handed, of Han ethnicity, had normal or corrected-to-normal vision, had no history of mental or physical illnesses, had no contraindications for MRI, and were not currently using any psychiatric medications.

To validate the performance of the decoders developed in the discovery cohort, we included an independent replication cohort comprising 35 healthy participants (18 females; mean ±SD age = 20.74 ±1.56 years) recruited from UESTC utilizing the same experimental paradigm as the discovery cohort. This replication cohort was collected as part of a separate study employing a double-blind design, and the data acquisition proposal and analyses were pre-registered before the start of the validation procedure. Each selected participant received a placebo nasal spray (consisting of glycerin, sodium chloride and purified water, without any pharmacological components) 45 minutes before the MRI scanning, and the administration of the placebo followed a standardized protocol.

For both the discovery and replication cohorts, written informed consent was obtained from all participants prior to their participation, and the study received approval from the Ethics Committee of UESTC, adhering to the latest revision of the Helsinki Declaration. Upon completion of the experiment, participants received a monetary compensation.

### Social exclusion and social separation stimuli

A total of 186 stimuli, including 27 exclusion videos, 29 inclusion videos, 40 separation videos, 30 company videos, and 60 control videos), each lasting between 10 and 15 seconds, were carefully chosen from the Internet. All video clips were uniformly adjusted to ensure consistency in resolution and size. The social exclusion stimuli feature individuals experiencing ostracism or being distanced by their peers, contrasting with the social inclusion stimuli, which showcase individuals engaging happily with their peers. Conversely, social separation stimuli depict individuals undergoing separations from their family members, romantic partners, and friends, whereas social company stimuli highlight individuals enjoying warm moments with their loved ones. Additionally, neutral stimuli present individuals calmly and quietly coexisting with strangers in the same space, without any overt social interaction.

An independent sample of 60 participants (30 females; mean±SD age=21.47±2.20 years) were recruited to screen the stimulus materials prior to the formal experiment. Each stimulus was categorized based on its valence (positive, negative or neutral) and subsequently evaluated using a 5-point Likert scale for arousal, intensity and pain empathy (‘how much do you feel pain towards the content of the video segment’). Following this screening process, a total of 120 stimuli were selected, including 22 for social exclusion, 22 for social inclusion, 22 for social separation, 22 for social company, and 32 control stimuli (half for the exclusion pain empathy task as the control condition, the remaining half for the separation pain empathy task).

Following the screening process, a total of 120 stimuli were selected, including 22 for social exclusion, 22 for social inclusion, 22 for social separation, 22 for social company, and 32 control stimuli (half for the exclusion pain empathy task as the control condition, the remaining half for the separation pain empathy task). An additional independent sample of 21 participants (10 females; mean±SD age=22.71±2.57 years) was recruited for behavioral ratings for the included 22 exclusion videos and 22 separation videos. They were required to categorize the stimuli as either social exclusion or social separation and evaluate the physiological distance between characters, as well as the familiarity ratings using a 5-point Likert scale. Based on the behavioral ratings, all participants were able to precisely discern whether the stimulus depicted social exclusion or social separation. Notably, 95% of the participants perceived the physiological distance between characters in the video clips featuring social exclusion as being considerable, and 86% considered the physiological distance in clips representing social separation to be relatively close. Additionally, there was no significant difference in participants’ familiarity levels between the two types of stimuli (**Supplementary Figure. 1**).

### Experimental paradigm

A total of 60 video stimuli (22-social exclusion; 22-social inclusion and 16-control) for social exclusion pain empathy task were randomly distributed across three runs, with each run containing 20 stimuli. Each trial started with an average 1s jitter (0.5-1.5s). Participants were instructed to watch the social experiences of characters in the videos, which lasted from 10 to 15 s, from a third-person perspective. Following a 3-5s jitter, they were asked to rate their level of pain empathy for each stimulus using a 5-point Likert scale (from ‘not at all’ to ‘strongest’, **see Figure 1A**) in 2s. The stimulus presentation was programmed with Psychopy-2022.1.3 (https://www.psychopy.org). The design for the social separation pain empathy task was identical to that of the social exclusion pain empathy. The order of these two tasks was counterbalanced across participants. Out of the scanner, all participants evaluated the arousal and intensity rating using five-point Likert scales (1= very low, 5= very high).

### MRI data acquisition and data preprocessing

The MRI data were acquired on a 3.0T GE MR750 system (General Electric Medical System, Milwaukee, WI) with an 8-channel phased array head coil. Functional MRI data were collected using T2*-weighted echo-planar imaging (EPI) pulse sequence with the following parameters: repetition time (TR) = 2000 ms, echo time (TE) = 30 ms, flip angle = 90°, field of view (FOV) = 240 mm × 240 mm, voxel size = 3.75 × 3.75 × 4 mm, resolution = 64 × 64, and 39 slices. High-resolution T1-weighted anatomical images were collected to enhance the normalization of functional images using a spoiled gradient-recalled (SPGR) sequence with the following parameters: TR = 6 ms, TE = 2 ms, flip angle = 9°, FOV = 256 × 256 mm, voxel size = 1 × 1 × 1 mm, 156 slices, and a slice thickness of 1 mm. All MRI data underwent preprocessing using the standard workflow in fMRIPrep 22.0.0.

The first five volumes of each run were discarded to achieve stabilization of image intensity. All MRI data underwent preprocessing using the standard workflow in fMRIPrep 22.0.0 ^66^, a process based on Nipype 1.8.3 ^67^. This workflow is an automated and flexible pre-processing pipeline that utilizes various neuroimaging software packages. The preprocessing steps include: correction of intensity non-uniformity in T1w images using the Advanced Normalization Tool (ANTs 2.3.3) ^68^ with the N4BiasFieldCorrection ^69^ algorithm, and skull-stripped through a template-based brain extraction procedure(template: OASIS30ANTS); brain tissue segmentation of cerebrospinal fluid(CSF), white matter, and gray matter in the brain-extracted T1w images using FSL’s FAST(FSL 6.0.5.1; https://fsl.fmrib.ox.ac.uk/fsl) ^70^; volume-based spatial normalization of T1w images to standard space (template: MNI152NLin2009cAsym) was conducted via nonlinear registration with ANTs by using brain-extracted versions of both the T1w images and the ICBM 152 Nonlinear Asymmetrical template version 2009c ^71,72^.

Functional data underwent slice-timing correction with AFNI’s 3dTshift ^73^ and motion corrected with FSL’s MCFLIRT ^74^. Subsequently, the participant’s corresponding T1w images were co-registered with six-degree-of-freedom registration via FSL’s FLIRT ^75^ based on boundary information ^76^. The forward deformation parameters obtained from the segmentation process were then applied for normalization to Montreal Neurological Institute (MNI) space (2×2×2 mm voxel size). Spatial smoothing was performed using an 8 mm full-width at half-maximum (FWHM) Gaussian kernel.

### General linear model (GLM) analysis

First-level general linear model (GLM) analysis was conducted on pre-processed fMRI data based on MATLAB (version 2020b, MathWorks), using the SPM12 software (https://www.fil.ion.ucl.ac.uk/spm/software/spm12/), and customized MATLAB code. This study involved three independent subject-level GLM analyses.

The first GLM model was employed to obtain subject-specific condition beta images and we included five separate boxcar regressors including Exclusion vs. Control, Inclusion vs. Control, Separation vs. Control, Company vs. Control, and pain rating period. The second GLM model adopted a parameter modeling approach, utilizing subject-level self-reported subjective pain empathy ratings on a 1-5 scale as labels during the presentation of video stimulus materials. This model aimed to capture neural activity changes across various stimulus materials for each participant based on the degree of their subjective empathy (most participants utilized the entire range of scores ‘1-5’ in their subjective pain empathy ratings, but some participants used only a subset of these scores). The third GLM model was established based on 120 videos (60 for exclusion pain empathy task and 60 for separation pain empathy task) to create single-trial images.

For each participant, the three GLM models concatenated data from a total of six runs, convolved with the classic hemodynamic response function (HRF) in SPM12, and applied a high-pass filter of 128s to remove low-frequency drift. Nuisance variables encompassed six estimated head motion parameters (X, Y, Z, roll, yaw, and pitch), as well as their squares, their derivatives, and the square of derivatives for each run (24 columns in total).

### Multivariate pattern analysis

The MVPA analysis aimed to precisely identify the weighted brain activity patterns specifically associated with pain empathy evoked by social exclusion and social separation. Here, linear support vector machines (SVMs) algorithm from Spider toolbox (http://people.kyb.tuebingen.mpg.de/spider) with parameters (C= 1, optimizer= andre) was applied to the data processed with first-level general linear model (GLM) in the discovery cohort.

The discovery cohort encompassed 260 contrast images, derived from 65 participants, each experiencing four conditions (Ex-Con, In-Con, Sep-Con, and Com-Con, i.e., 65×4). To eliminate general neural activation triggered by viewing video stimuli, such as sensory, visual, and emotion-related activations ^77,78^, the four conditions were obtained by subtracting their respective control conditions, signifying exclusion, inclusion, separation, and company, respectively ^79^. The following four pattern classifiers were developed: PSSE, PSSS, PSSEvS, and PSSEaS. Utilizing the one-against-all approach ^43,80^, we trained the PSSE model to specifically discriminate between exclusion and the remaining three conditions. Similarly, the PSSS model was trained to classify between separation and the other three conditions. Furthermore, the PSSEvS model was designed to capture the stimuli-specific differences between exclusion and separation, while the PSSEaS model aimed to explore the common between exclusion and separation.

### Evaluation the performance of models

To rigorously evaluate the pattern performance of both the PSSE and PSSS pattern classifiers, we employed a robust 1000 iterations of a 10×10-fold cross-validation procedure ^81,82^. Utilizing MATLAB’s cvpartition function ^83^, the results of the dot product between the vectorized univariate general linear model (GLM)-derived fMRI activation maps and the weight maps of PSSE and PSSS were evenly allocated across ten sub-samples. According to the data from the nine sub-samples as a training dataset, we computed the optimal hyperplane using MATLAB’s fitcsvm function ^84^. Subsequently, performance evaluation was conducted on the remaining one sub-sample (testing set). The entire process was repeated ten times, with each iteration ensuring that every sub-sample served as a training dataset and as a testing dataset. This iterative approach enhances the robustness of our evaluation, as it allows for a comprehensive assessment of the pattern classifiers’ performance across various training-testing splits. Additionally, the entire performance evaluation process was replicated 1000 times, with each iteration involving a unique partitioning of the 10 sub-samples. This extensive repetition not only helps avoid potential biases associated with specific training-testing splits ^45,85^, but also provides a reliable and convergent estimate of the pattern performance ^41^. For both PSSEvS and PSSEaS, the classification performance was computed using the roc_plot function from the CanlabCore tools (https://github.com/canlab/CanlabCore).

### Identifying core systems involved in social pain empathy

The core brain regions involved in social pain empathy are identified as those voxels that consistently contribute to model classification (i.e., model weights) and exhibit an interpretable relationship with the model classification process (i.e., structure coefficient) ^39,41^. Previous research posits that both model weights and structural coefficients are essential for model interpretation ^86,87^. Model weights represent the classification coefficients and the directional effects of variables, controlling for the influence of other variables in the model. In contrast, structural coefficients indicate the direction of the relationship between variables and the model’s outcome without controlling for other variables ^45^. The specific steps we conducted are as follows:

Step 1: Generation of model weight maps. To identify the cerebral regions that consistently contributed to the classification, we employed a bootstrap test. This involved 10,000 bootstrap samples (with replacement) from the discovery cohort and performed SVM on each sample set. Subsequently, we computed Z-scores and two-tailed uncorrected P-values for each voxel, based on the mean and standard deviation obtained from the bootstrap distribution. Based on the corresponding P-values, we applied a threshold to the statistical map after correction for multiple comparisons (FDR q<0.05, **Supplementary Figure. 2A**).

Step 2: Generation of model encoding maps. Considering that the core regions are likely associated with the empathy process and to minimize noise in the data, the Haufe transformation (https://github.com/kabush/kablab) was employed to transform the weight map obtained in Step 1 (i.e., PSSE, PSSS, etc., essentially, the backward models) into “structural coefficients” (i.e., forward models) ^88^. These coefficients establish a direct mapping between each voxel and the response (fitted values) of the multivariate models. By conducting a single-sample t-test with FDR correction (q < 0.05), we identified significantly activated brain regions that not only demonstrated reliable classification of empathy but also exhibited a correlation with the empathy process (**Supplementary Figure. 2B**).

Step 3: Identification of core regions. The core regions were identified as those voxels that consistently exhibited a correlation and predictive for the target outcome (i.e., subjective social pain empathy). They were determined by their consistent performance in both backward and forward models ^54,89,90^(**Supplementary Figure. 3**).

### Spatial similarity analysis between model encoding maps

The entire brain cortex and subcortical regions, based on the 246-region version of the Brainnetome Atlas, were divided into 7 networks and 24 distinct brain regions. Spatial similarity was quantified using the cosine similarity metric, comparing the activation patterns within the core regions identified through both the PSSE and PSSS models with each individual regional interest (ROI) or network. By employing river plots, we were able to assess the degree of overlap and consistency between the activation patterns associated with social pain empathy and the anatomical parcellations defined by the Brainnetome Atlas ^39,50^. All voxels were maintained as positive values for interpretation.

### Extended exploration of decoder performance

Here, we integrated and evaluated previously developed whole-brain pain decoders from prior studies: the Facial Expressions Pain Experience (FEPain), decoding the pain evoked by observing others’ painful facial expressions), the Noxious Stimulation Pain Experience decoder (NSPain), decoding the pain induced by observation of noxious stimulation of others’ body limbs, and the Physiological Pain Empathy Signature (PHPain), a combined model between FEPain and NSPain ^46^; the Social Rejection Experience decoder (REPain), which decodes observing participants’ own ex-partner picture as rejection pattern) ^43^, the Physiological Heat Pain decoder (HEPain) ^43^ and the Stimulus Intensity Independent pain signature-1 (SIIPS1) ^53^, induced by heat pain. These decoders were integrated to further validate and explore the decoders we developed in the current study. Specifically, we first applied the aforementioned decoders to our datasets including the discovery cohort (n=65) and replication cohort (n=35). Subsequently, we evaluated our decoder’s performance on independent validation cohort 1 and validation cohort 2, the original dataset utilized for developing NSPain and FEPain (n=238, 118 females; mean ±SD age=21.58 ± 2.32 years) ^46^, the generalization cohort 1, which comprised participants recalling the episodic images of others’ bodily pain stimuli ranging from blurry to clear (n=77, 40 females, mean age ±SD: 20.13±2.39 years), and the generalization cohort 2, consisting of visually evoked subjective disgust (n=78, 16 females, mean age ±SD: 21.13±2.18 years).

### Multilevel two-path mediation analysis

To further investigate the intricacies of the social pain empathy neural decoder developed in this study, we employed a multilevel two-path mediation analysis using the Mediation Toolbox (https://github.com/canlab/MediationToolbox) ^42^. This approach aims to examine whether the covariance between the independent variable X and the dependent variable Y (i.e., empathic responses to social stimuli) can be attributed to the mediating variable M.

Within the framework of multilevel two-path mediation ^42,53^, X-M, M-Y, and the two X-Y relationships before and after mediation are defined as path a, path b, path c, and path c’, respectively. Specifically, the difference between the total (c) and direct effect (c’) of X-Y (i.e., c − c’) can be determined by testing the significance of the product a×b, which represents the multiplication of the path coefficients of path a and path b. If all three components (i.e. a, b, and the product a × b) are significant, the mediation effect is considered significant. In this study, we conducted two mediation analyses: (1) whether the neural decoder for physiological pain empathy serves as a mediator in the relationship between the social pain empathy decoder and responses to social empathy stimuli; (2) whether the neural decoder for personal experiences of separation mediates the relationship between the social pain empathy decoder and responses to social empathy stimuli. Here, the dot multiplication of the mentioned decoders with single-trial contrast images (each participant included 60 exclusion class contrast images and 60 separation class contrast images) was performed, and the statistical significance of the mediation effects was assessed using bootstrap tests with 10,000 iterations.

## Acknowledgements

This work was supported by the Natural Science Foundation of Sichuan Province [grant number 2022NSFSC1375 – WHZ; 2023NSFSC1185 - XQZ], National Natural Science Foundation of China (NSFC) [grant number 31530032 - KMK] and Key Scientific and Technological projects of Guangdong Province [grant number 2018B030335001 - KMK]. Disclaimer: Any opinions, findings, conclusions or recommendations expressed in this publication do not reflect the views of the Government of the Hong Kong Special Administrative Region or the Innovation and Technology Commission.

## Authorship Contribution Statement

X.Z., and W.Z. designed the experiment, analyzed the data and drafted the manuscript. X.Z., P.Q., Q.L., C.L., L.L., X.G., K.F., C.L., and X.Z. conducted the experiment. K.M.K. and B.B. provided feedback and revised the manuscript.

## Declaration of Competing Statement

The authors declare no competing interest.

## Notes

### Competing Interest Statement

The authors have declared no competing interest.

